# Controllable Point-Light Displays Implemented in a game engine for biological motion research

**DOI:** 10.64898/2026.01.05.697634

**Authors:** Milad Yekani, Shirin Jamshidi, Babak Feyzifard, Abdol-hossein Vahabie

## Abstract

Biological motion perception plays a critical role in survival and social communication across species. Point-light displays (PLDs), which represent body movement using only a small set of joint markers, have long served as an effective tool for isolating motion cues from other visual features. However, existing methods for generating PLDs, ranging from filmed actors with reflective markers to markerless motion extraction and motion-capture datasets, present limitations in cost, accessibility, flexibility, or ecological validity. In particular, many laboratories lack the resources to create customizable stimuli that allow systematic manipulation of movement parameters. In this article, we introduce a practical and easily modifiable method for producing fully controllable 3D PLDs using freely available animation rigs and the Unity game engine. Our approach enables real-time control of depth cues and directional motion without the need for motion capture equipment or specialized filming environments. To demonstrate the utility of the method, we conducted a psychophysical experiment comparing the perception of biological and non-biological motion. The results replicate the well-documented forward-motion perceptual bias for biological stimuli and highlight differences in how observers interpret motion direction across stimulus classes. This method offers a convenient, accessible, and adaptable tool for research on motion perception and social cognition.

## Introduction

body motion is an essential ability for species to escape predators, pursue prey, and communicate socially. Our brains perceive biological motion as distinct stimuli, different from other types of motion ^12^. The presence of specialized neural mechanisms for biological motion in diverse species, such as humans ^3^ and birds ^4^, demonstrates its ecological importance and makes it an attractive target for psychological and neuroscientific research. Since the 1970s, point-light displays have become a foundational tool for studying biological motion perception. First introduced by Johansson (1973), this method uses illuminated points placed on key joints to isolate motion cues while removing other visual details. Creating such stimuli remains a critical step in research on body-motion perception, and multiple techniques have been developed, each with advantages and limitations. This article introduces a new method for generating 3D human motion and demonstrates its effectiveness by producing fully controllable point-light walker stimuli.

A straightforward way to create point-light displays is by recording actors wearing retroreflective markers or lights attached to their joints. Johansson’s original filming method is conceptually simple, but the resulting stimuli are difficult to modify parametrically (e.g., speed, limb phase, viewpoint), require specialized filming conditions, and do not preserve 3D structure for interactive graphical environments^5,6^. As a result, these stimuli offer limited capacity for manipulating depth cues or systematically altering motion information in perception experiments.

An alternative is using **markerless pose estimation**, where joint coordinates are inferred from video using machine learning. Recent deep-learning-based systems such as **OpenPose, DeepLabCut**, and **MediaPipe Pose** have dramatically improved motion extraction precision ^7–9^. However, these systems are computationally demanding and remain imperfect: small tracking errors can lead to jitter and degraded perceptual naturalism in the resulting stimuli, limiting ecological validity in psychophysical tasks.

Motion capture studios using reflective markers and infrared multi-camera setups provide highly accurate 3D joint trajectories and are widely used in animation and biomechanics research. Several publicly available datasets allow researchers to generate PLDs without collecting new data. However, these datasets offer **fixed motions**, which limits their usefulness for experiments requiring systematic manipulation—for example, altering emotional expression or social intention conveyed by motion^11,12^. While **IMU-based** systems are a portable and scalable solution for recording kinematic information^13^, they remain relatively expensive, require calibration, and are typically used in applied fields such as gait rehabilitation rather than perceptual research.

Overall, current methods pose practical and methodological challenges for laboratories interested specifically in **perceptual** aspects of biological motion. Many labs are unwilling or unable to invest in high-cost motion capture systems, and available datasets often lack flexibility for controlled manipulation. There is thus a strong need for a **low-cost, modifiable, and high-precision** method for generating point-light stimuli that preserves ecological motion structure while enabling fine experimental control.

## Methods

### Creating stimuli

#### 1) PLD Avatar

We downloaded FBX files of a walking character (named Ybot) from the Mixamo website. We just downloaded the rig without the mesh. Then, spheres were attached to each main joint in the Blender software environment. We can export the FBX file at this stage, which is several dots attached to each joint with a walking animation. This FBX file can be used as an avatar in the Unity software.

#### 2) Non-biological Movement Stimuli

In this study, a Unity environment was developed to generate non-biological stimuli by creating 500 spheres that move along one axis, with customizable proportions moving either towards or away from the observer (camera). The goal was to simulate varying directional movements in 3D space, mimicking non-biological motion patterns with controlled proportions of approach and retreat. We created 11 stimuli corresponding to 11 degrees of manipulation on the perspective cue, which included (15, 22, 29, 36, 43, 50, 57, 64, 71,78, 92) percent of dots moving toward the observer.

**Figure 1.**
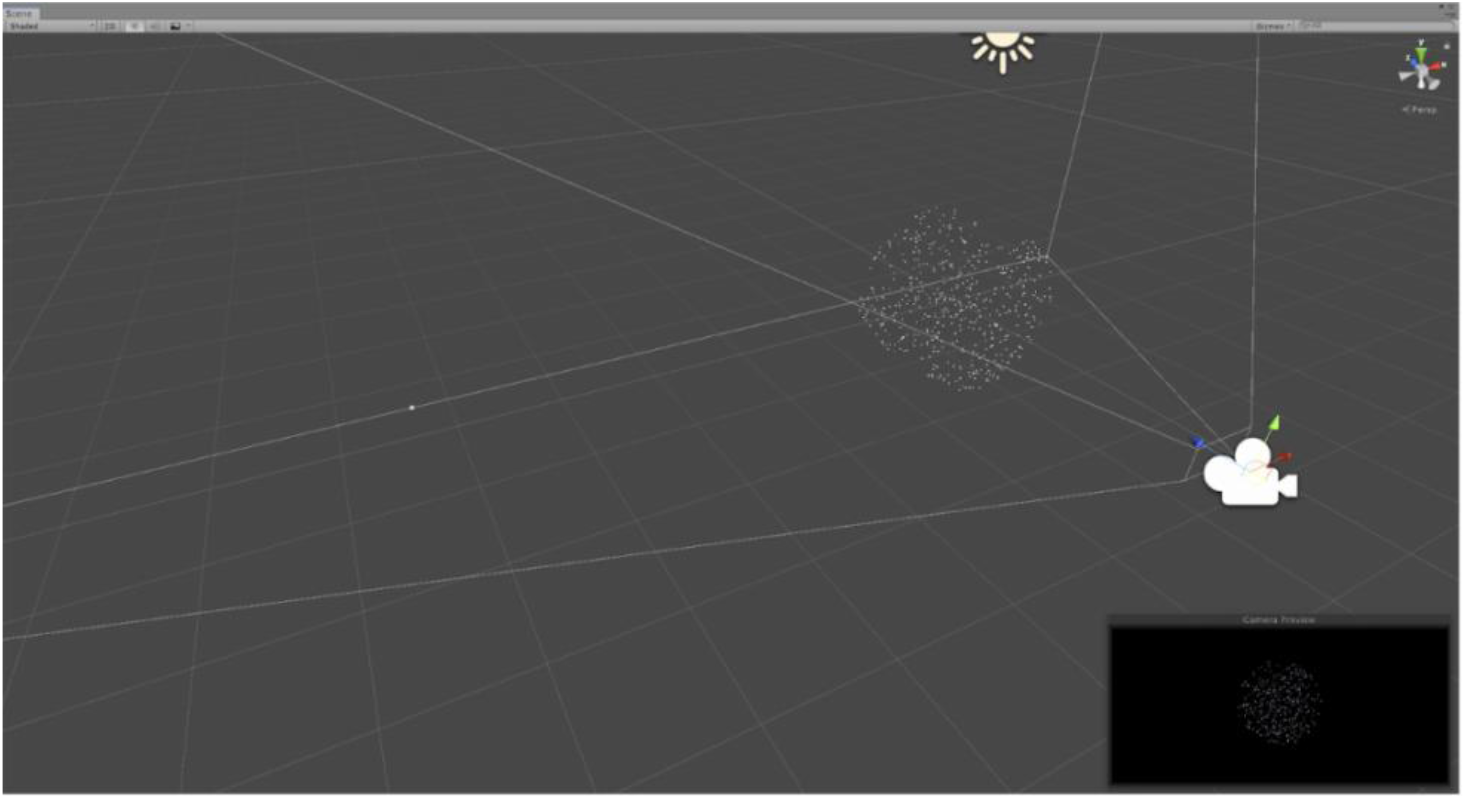
Non-biological movements are produced by 500 randomly generated spheres with different proportions of moving-toward or moving-away spheres. The bigger window shows the scene view of the Unity environment, and the black window in the corner shows what is rendered from the camera’s perspective.

### Creating the Unity environment

We programmed individual dots in Unity to mimic precisely the two coordinates on the vertical plane parallel to the character’s body. The depth coordinates of the points were assigned to spheres after they were multiplied by an adjustable parameter (named Z coefficient). The camera was in front of the avatar (at 4 unit distance). We used a specific rendering setting to avoid the detectability of light direction on spheres. Negative values of the Z parameter cause the PLD to move away from the camera, and positive values work oppositely. We used a black background and colored dots. The camera moves in the depth dimension to keep the fixed distance between it and the avatar’s base bone. This prevents the avatar’s picture from getting smaller when it moves away and bigger when it comes toward it. (Refer to the available codes and documents here). We created our stimuli by filming the avatar through the camera and separately for each different z parameter. We used 11 levels of different z parameters (-1,-0.8, -0.6, -0.4, -0.2, 0, 0.2, 0.4, 0.6, 0.8, 1).

**Figure 2.**
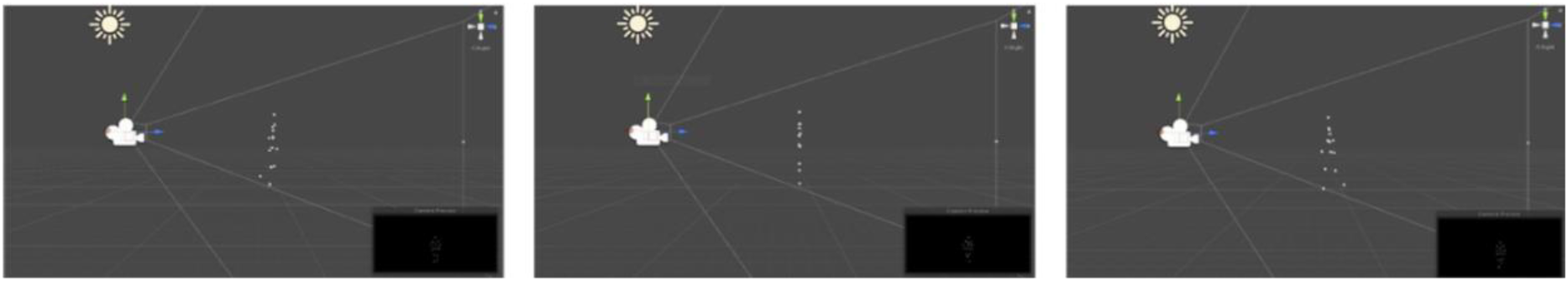
Applying varying degrees of perspective cues (left:-0.8, middle: 0, right:0.6) on the PLD avatar in Unity’s environment. The larger window displays the scene view of the Unity environment, while the black windows in the corners show what is rendered from the camera’s perspective.

### Task design

In this study, participants watched two sets of videos to see how they perceive movement. The first set showed biological movement with walking characters. After a 10-minute break, participants watched the second set of non-biological movement videos, which displayed dots forming circles. Each set had 220 videos, arranged randomly to keep the same order for everyone. The videos were split into 20 rounds, with 11 different difficulty levels in each round. During the videos, we recorded participants’ answers and the time it took them to respond. In the biological movement videos, participants answered “Forward” if they saw the character moving forward and “Backward” if they thought it was moving backward. In the non-biological videos, “Forward” meant the dots appeared to be moving closer, while “Backward” meant the dots seemed to be moving away. We measured reaction times from when the video started until the participant pressed a key. They used the down arrow key for “Forward” and the up arrow key for “Backward.”Each video was shown for up to five seconds. Participants were told to respond within this time, but could answer as soon as they noticed the movement direction. If they didn’t respond in time, the video would disappear briefly, and missed videos were replayed at the end of the set. Participants were encouraged to respond to as many videos as possible.

**Figure 3.**
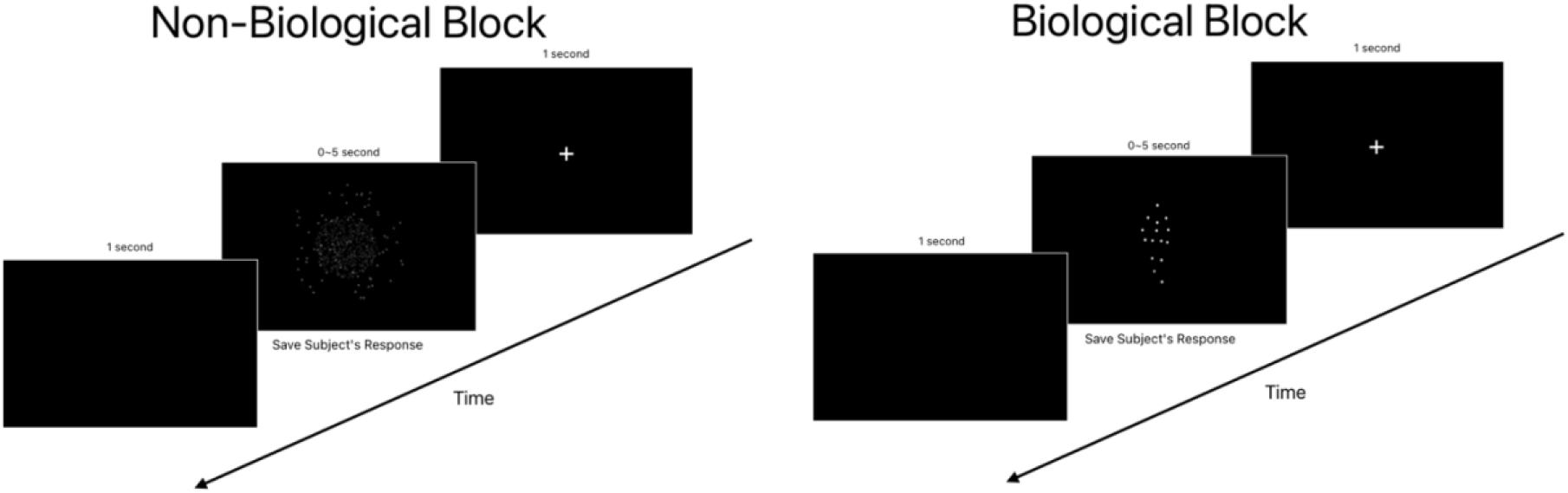
A block of non-biological and biological diagrams. Each block contains a black screen with a white “Plus” sign, which is exposed for a second to the subject. The next scene of the block is the designed video, which has a time limit of 5 seconds, but it could take less if the subject responds faster than 5 seconds. Lastly, a fully black screen is shown for a second.

### Data analysis

All participants’ responses were analyzed, and the number of forward responses for each label was calculated. This data was then plotted on a graph to visually visualize the pattern of participants’ choices for biotic and abiotic stimuli. This initial step helped us to get a general idea of how responses were distributed and individuals’ initial orientations toward each stimulus category. In the next step, the Boltzmann curve was used to model the choices, which are more accurate. This curve is one of the most widely used tools in psychometric data analysis and cognitive neuroscience, used to model binary choices (e.g., choosing between two options). This curve uses a logistic function to model the probability of choosing a particular option as a function of differences in stimulus characteristics (such as value, intensity, or biological/non-biological characteristics). In this study, we used the Boltzmann curve to examine participants’ biases toward biological and non-biological stimuli.

To calculate the bias, the midpoint of the Boltzmann curve was taken as the criterion. This value indicates at what value of feature difference the participant was equally hesitant between the two options. If this point is far from zero, it indicates the presence of a bias or unwanted tendency towards the biological or abiotic stimulus. The amount of this shift (bias) was calculated for each individual and used in the final analyses. The bias occurs when the probability of choice is 0.5, and at this point, the participant is least confident about their perception, which can vary for each individual.

## Results

**Figure 4.**
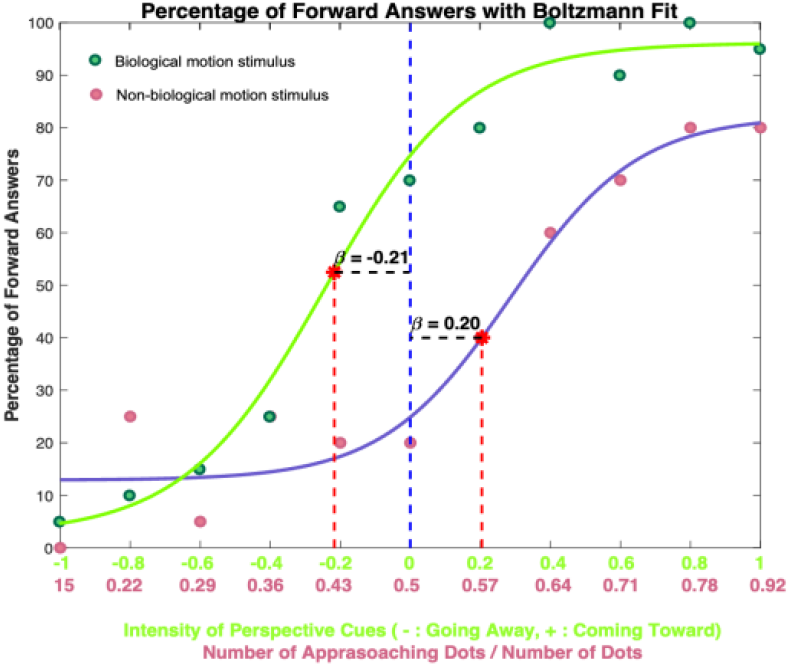
This figure belongs to subject 11, showing the data points for biological (green points) and non-biological (pink points). We fit a Boltzmann curve for biological data points (green curve) and non-biological data points (purple curve). Mean points for each curve have been written and named Beta, which are their distance from the mean point of the labels. The horizontal and vertical indicate the video labels for biological (green labels) and non-biological (pink labels) videos.

**Figure 5.**
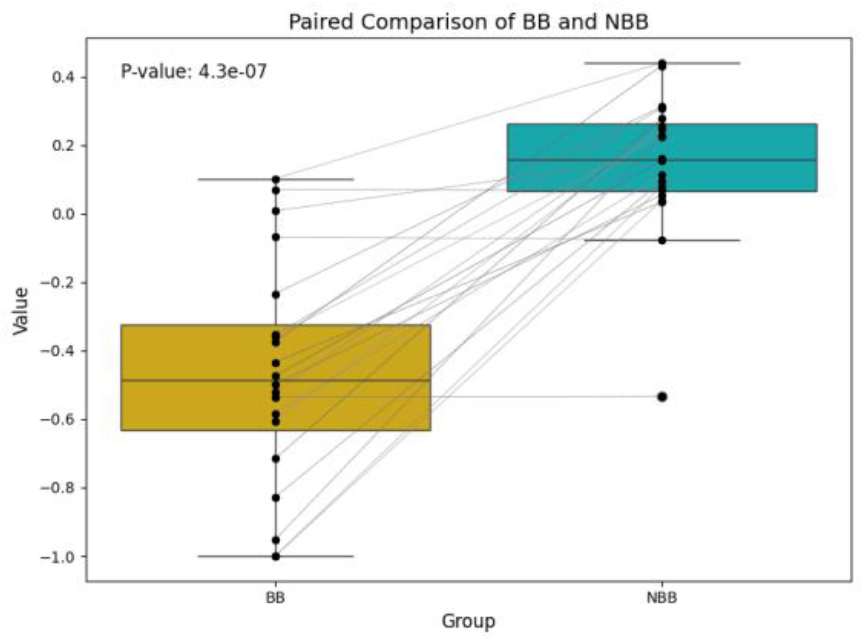
This figure indicates the biases for biological (BB) and non-biological (NBB) data points for each subject. The corresponding biases for each subject are connected with a gray line. The mean bias and standard deviation for biological data points are -0.468 and 0.326, respectively. These values for non-biological data points are 0.146 and 0.204.

## Psychometric curve

The figure on the left contains the answers to subject 11 of the test. The vertical axis shows the percentage of forward answers that the subject has selected. The horizontal axis shows the labels of videos, which are different for biological and non-biological videos due to their different nature.

Each green circle shows the ratio of the videos that the subject selected the forward answer to the total number of videos for a label, in percentage for biological videos, and each pink circle shows the corresponding percentage for non-biological videos. Then we fit a Boltzmann curve for each set of biological and non-biological data points, and then we found the mean point of these curves by measuring the minimum and maximum percentage in each curve and dividing the summation of the maximum and minimum by two, and named them Beta.

We can see that participants are more likely to perceive biological stimuli as more forthcoming. Beta value for biological bias for all participants is -0.468, and the corresponding value for non-biological bias is 0.146. We can also see that participants are more likely to perceive the nonbiological motion stimuli as moving away from them. Standard deviation for the biological biases is 0.326, and this value for non-biological biases is 0.204.

These results can be seen in each subject’s responses, similar to the presented sample–the left figure. For this subject, the Beta value for the biological Boltzmann curve is -0.21, which means it is 0.21 behind the center of the labels, which is label 0. This value for the non-biological Boltzmann curve is +0.20, which means it is 0.20 ahead of the center of the labels, which is label 0.5.

## Discussion

This article presents a new and convenient technique for creating 3D point-light displays (PLDs) to analyze both biological and non-biological movement. While traditional methods are effective, they come with challenges related to cost, adaptability, and accessibility^1,2^. Our goal was to address these limitations by devising an easily applicable solution for producing PLDs in a 3D visual environment. Our psychophysical study demonstrated the efficacy of this approach, illustrating its utility for investigating perceptual biases and motion-direction interpretation.

The proposed method offers significant advantages over traditional techniques such as filming actors wearing retroreflective markers, markerless motion capture, or using pre-existing motion capture datasets. Unlike the setup-dependent filming methods initially introduced by Johansson (1973), our method allows researchers to generate stimuli entirely within a computer graphics environment, removing the need for specialized equipment, motion capture studios, or actor performances. Moreover, the system can be modified in real time, enabling flexible experimental manipulations such as altering depth cues or movement kinematics ^3,14^. Pre-existing motion datasets are often difficult to adapt for experiments requiring emotion modulation, action intention cues, or social signaling ^11,12^

. Our method overcomes this by allowing parameterized control over motion.

Our findings also reinforce well-established perceptual biases in biological motion interpretation. Participants exhibited a strong forward-direction bias, with ambiguous movements more often perceived as approaching, consistent with prior research suggesting that biological motion perception is tuned toward detecting potential social agents or approaching organisms ^15–17^. The more accurate interpretation of non-biological motion based on perspective cues indicates distinct cognitive mechanisms for interpreting biological versus abstract motion patterns^18,19^.

This method is particularly advantageous for labs with limited funding or without access to motion capture systems. While deep-learning-based markerless motion capture systems (e.g., OpenPose, DeepLabCut) have recently become more accessible, they remain computationally intensive and can introduce jitter or instability, thereby reducing ecological validity ^8,20^. Our approach produces stable, repeatable, and customizable outputs without these drawbacks. Previously, researchers have used ordinary game characters in VR settings to study how movement affects human behavior ^21^. The PLD humanoid avatars used in this study can easily replace them to observe the pure effect of bodily motion in VR environments.

The versatility of this method allows it to be adapted for numerous perceptual research applications. For example, researchers could systematically manipulate gait patterns or expressiveness to study emotional recognition from movement ^12^, or examine clinical populations such as autism spectrum conditions, where biological motion processing can be atypical ^22,23^. Future research could integrate neuroimaging to investigate neural correlates of these perceptual biases ^24^, or extend the method to multi-agent interaction contexts.

## Conclusion

This article presents a new, adaptable, and convenient technique for creating 3D point-light displays for perceptual research. Our psychophysical experiment illustrated the effectiveness and feasibility of the method, indicating its ability to faithfully replicate biological motion and uncover significant perceptual biases, such as the forward direction bias in biological motion. This approach tackles the main drawbacks of current methods, equipping researchers with an affordable, customizable tool for investigating motion perception across a wide range of fields, from fundamental research to clinical applications. By providing a simple way to produce point light stimuli, our method enables researchers to develop and conduct personalized experiments that were previously limited by the complexity and rigidity of traditional motion capture systems. This innovative approach bridges the accessibility and experimental precision gap, rendering it a valuable instrument for enhancing the exploration of biological motion perception.

